# A piggyBac-based toolkit for inducible genome editing in mammalian cells

**DOI:** 10.1101/448803

**Authors:** Megan D. Schertzer, Eliza Thulson, Keean C.A. Braceros, David M. Lee, Emma R. Hinkle, Ryan M. Murphy, Susan O. Kim, Eva C.M. Vitucci, J. Mauro Calabrese

**Author notes:** National Institute for Environmental Health Sciences, Research Triangle Park, NC 27709.

## Abstract

We describe the development and application of a novel series of vectors that facilitate CRISPR-Cas9-mediated genome editing in mammalian cells, which we call CRISPR-Bac. CRISPR-Bac leverages the piggyBac transposon to randomly insert CRISPR-Cas9 components into mammalian genomes. In CRISPR-Bac, a single piggyBac cargo vector containing a doxycycline-inducible Cas9 or catalytically-dead Cas9 (dCas9) variant and a gene conferring resistance to Hygromycin B is co-transfected with a plasmid expressing the piggyBac transposase. A second cargo vector, expressing a single-guide RNA (sgRNA) of interest, the reverse-tetracycline TransActivator (rtTA), and a gene conferring resistance to G418, is also cotransfected. Subsequent selection on Hygromycin B and G418 generates polyclonal cell populations that stably express Cas9, rtTA, and the sgRNA(s) of interest. Using *Mus musculus*-derived embryonic and trophoblast stem cells, we show that CRISPR-Bac can be used to knockdown proteins of interest, to create targeted genetic deletions with high efficiency, and to activate or repress transcription of protein-coding genes and an imprinted long noncoding RNA. The ratio of sgRNA-to-Cas9-to-transposase can be adjusted in transfections to alter the average number of cargo insertions into the genome. sgRNAs targeting multiple genes can be inserted in a single transfection. CRISPR-Bac is a versatile platform for genome editing that simplifies the generation of mammalian cells that stably express the CRISPR-Cas9 machinery.

## Introduction

Within the last decade, the CRISPR (clustered regularly interspaced short palindromic repeat) bacterial immune system has provided researchers with multiple new methods to control gene expression in mammalian genomes. Co-expression of the Cas9 (CRISPR-associated protein 9) nuclease from *Streoptococcus pyogenes* along with an engineered single guide RNA (sgRNA) that targets a protein-coding exon is an effective way to introduce frameshift mutations in proteins of interest, owing to the fact that repair of the DNA break introduced by Cas9 often results in small deletions surrounding the cut site. Co-expression of Cas9 and multiple sgRNAs can also be used to excise larger regions from genes of interest, or to excise DNA regulatory elements. Expression of a catalytically-dead Cas9 (dCas9) fused to a transcriptional activation or repression domain can be used to up- or down-regulate gene expression when sgRNAs are targeted to promoters or regulatory elements of interest (Hsu et al., 2014; Wright et al., 2016).

Owing to the broad utility of CRISPR, multiple methods have been developed to deliver the CRISPR-Cas9 machinery to mammalian cells. Transient transfection of Cas9- and sgRNA-expressing plasmids, or of Cas9 protein and *in vitro* synthesized sgRNAs, are useful when the efficiency of transfection for the cell type of interest is high, and when the desired endpoint can be reached via transient expression of Cas9 and the sgRNA. Lentiviral delivery of Cas9/sgRNA vectors is also possible, and provides distinct advantages when transfection efficiency is low, or when the desired endpoint requires stable expression and or integration of Cas9/sgRNAs into the genome, such as for studies performed *in vivo* or for genome-wide phenotypic screens (Hartenian and Doench, 2015; Joung et al., 2017). However, delivery of the CRISPR machinery via lentivirus requires additional hands-on time, expertise, safety precautions, and cost relative to delivery via transient transfection.

The piggyBac transposon is a broadly used tool that allows DNA cargos up to 100 kilobases in length to be inserted into “AATT” sequences that are preferentially located in euchromatic regions of mammalian genomes (Cadinanos and Bradley, 2007; Ding et al., 2005; Li et al., 2011; Wang et al., 2008; Wilson et al., 2007). Owing to its high efficiency of transposition, piggyBac has been used in a wide range of applications, including in the stable expression of multi-subunit protein complexes, in the generation of transgenic mice and induced pluripotent stem cells, and in the large-scale production of recombinant proteins (Ding et al., 2005; Kahlig et al., 2010; Kaji et al., 2009; Li et al., 2013; Yusa et al., 2009).

Herein, we describe the creation and validation of a piggyBac-based system for inducible editing of mammalian genomes by CRISPR-Cas9. In the system, which we call “CRISPR-Bac”, two separate piggyBac cargo vectors, one that expresses an inducible Cas9 or dCas9 variant, and another that expresses an sgRNA and the reverse-tetracycline transactivator [rtTA; (Gossen et al., 1995)], are transfected into cells along with a plasmid that expresses the piggyBac transposase. A short period of selection is used to obtain cells that stably express both the Cas9 and sgRNA cargo vectors. We show that CRISPR-Bac provides a simple way to rapidly insert the CRISPR-Cas9 machinery into mammalian genomes to knockdown proteins, delete kilobase-sized genomic regions, and activate or repress transcription of protein coding genes and long noncoding RNAs (lncRNAs), without the additional cost and labor involved in the packaging and delivery of lentiviral particles to cells.

## Results

### Cloning of CRISPR-Bac

We modeled CRISPR-Bac (Figure 1) after the pX330 plasmid system in which a humanized version of the Cas9 enzyme from *Streoptococcus pyogenes* is co-expressed with a chimeric sgRNA driven by a U6 promoter (Cong et al., 2013). We cloned the Cas9 from pX330 into a doxycycline-inducible expression cassette in a piggyBac cargo vector that also expresses a gene conferring resistance to Hygromycin B. We then converted the dual BbsI sites in pX330, which are used to clone the sgRNA targeting sequence into that vector, into dual BsmbI sites. Like BbsI, BsmbI is a Type IIS restriction enzyme, and it generates overhanging ends that are identical to those generated by BbsI. We cloned the BsmbI-modified sgRNA expression cassette into a piggyBac cargo from (Kirk et al., 2018) that expresses a bi-cistronic message which encodes the rtTA*3* gene and a gene conferring resistance to G418 (originally cloned from Addgene plasmid #25735; (Shin et al., 2006)). The conversion of the pX330 BbsI sites, which are not unique in the rtTA-expressing vector, to BsmbI sites, allows the exact sgRNA design and cloning protocol for pX330 (Cong et al., 2013) to be used to clone sgRNAs into CRISPRBac.

**Figure 1.**
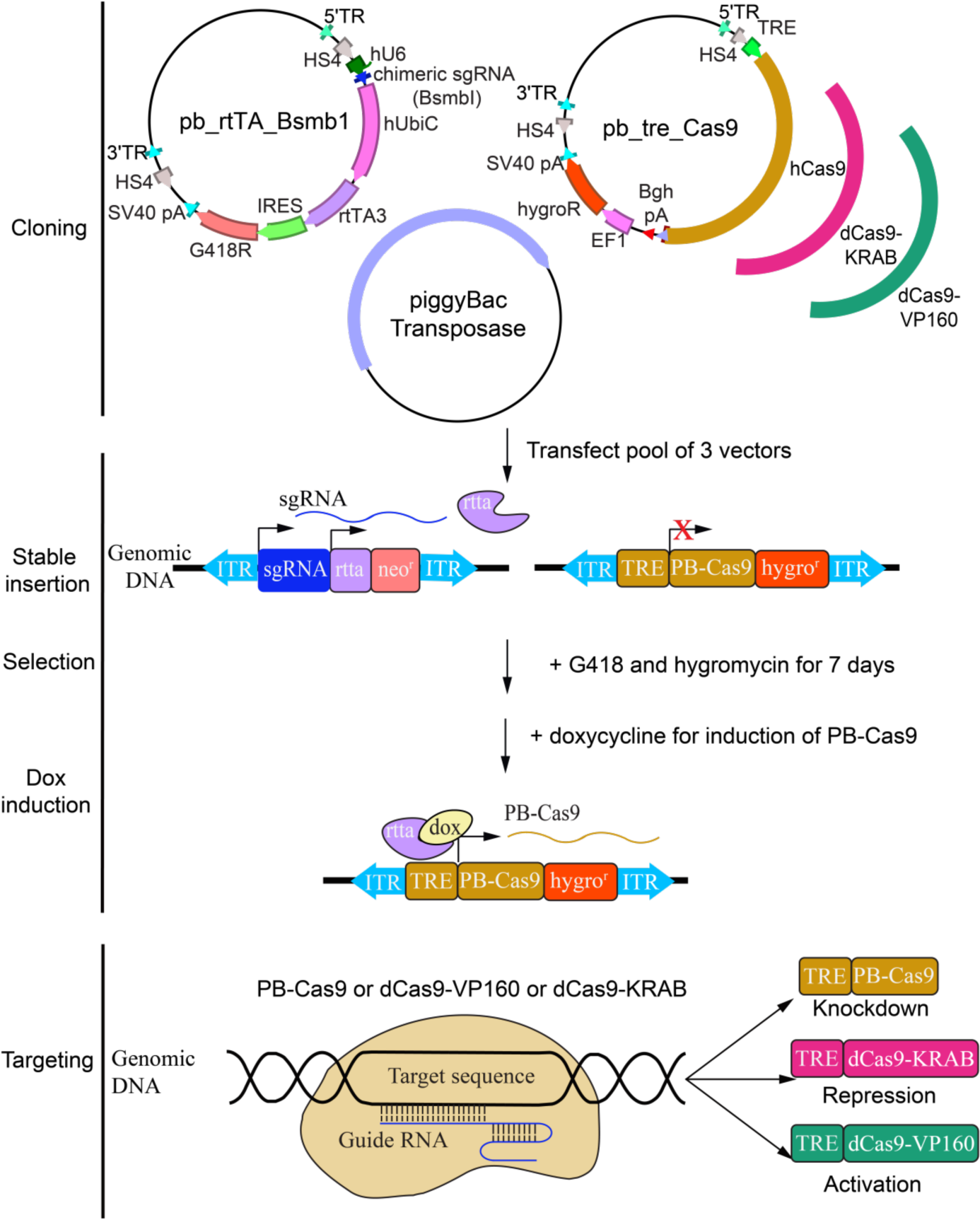
Experimental pipeline used for CRISPR-Bac. In a CRISPR-Bac experiment, a Cas9-expressing piggyBac cargo vector (or dCas9 variant) is co-transfected with a sgRNA- and rtTA-expressing piggyBac cargo vector and with a piggyBac transposase plasmid. Growth in Hygromycin B and G418 for 7 to 12 days selects for a population of cells that stably express an sgRNA of interest, and inducibly express a Cas9 or dCas9 variant. TR, piggyBac inverted terminal repeat. HS4, chicken bglobin insulator element. TRE, tetracycline responsive element (i.e. doxycycline inducible promoter). hCas9, dCas-VP160, dCas-KRAB from (Cheng et al., 2013; Cong et al., 2013; Kearns et al., 2014). EF1, EF1a promoter. HygroR, Hygromycin B resistance gene. SV40pA, SV40 polyadenylation signal. hU6-chimeric sgRNA from (Cong et al., 2013) with BsmbI sites replacing BbsI sites. hUbiC-rtTA3-IRES-G418 cassette was from pSLIKNeo (Shin et al., 2006).

### Knockdown of a protein-coding gene using CRISPR-Bac

We tested whether CRISPR-Bac could be used to knockdown a protein of interest in mouse embryonic stem cells (ESCs). We designed three sgRNAs targeting different exons of the *Ezh2* gene and cloned them into our sgRNA-rtTA-expressing vector using the protocol outlined in (Cong et al., 2013). We then co-transfected our inducible Cas9-expressing piggyBac vector, a plasmid expressing the piggyBac transposase, and either each sgRNA-expressing vector separately, or a pool of all three sgRNAs into ESCs. As a control, we transfected an sgRNA-rtTA-expressing vector into which we did not clone a specific sgRNA-targeting sequence (our “non-targeting sgRNA” control). After selecting ESCs on Hygromycin B and G418 for 10 days, we removed the selection drugs and added 1µg/ml of doxycycline to the media for four days to induce the expression of Cas9. To assess the extent of EZH2 knockdown, we performed western blot and immunofluorescence. Relative to the control ESCs, we observed greater than 60% reduction in EZH2 protein levels in the three lines expressing individual sgRNAs and more than 90% loss in the line expressing the pool of sgRNAs (Figure 2A, B). Immunofluorescence to EZH2 confirmed our western blot analysis (Figure 2C). These results demonstrate that CRISPR-Bac can be used to inducibly knockdown a protein-coding gene of interest.

**Figure 2.**
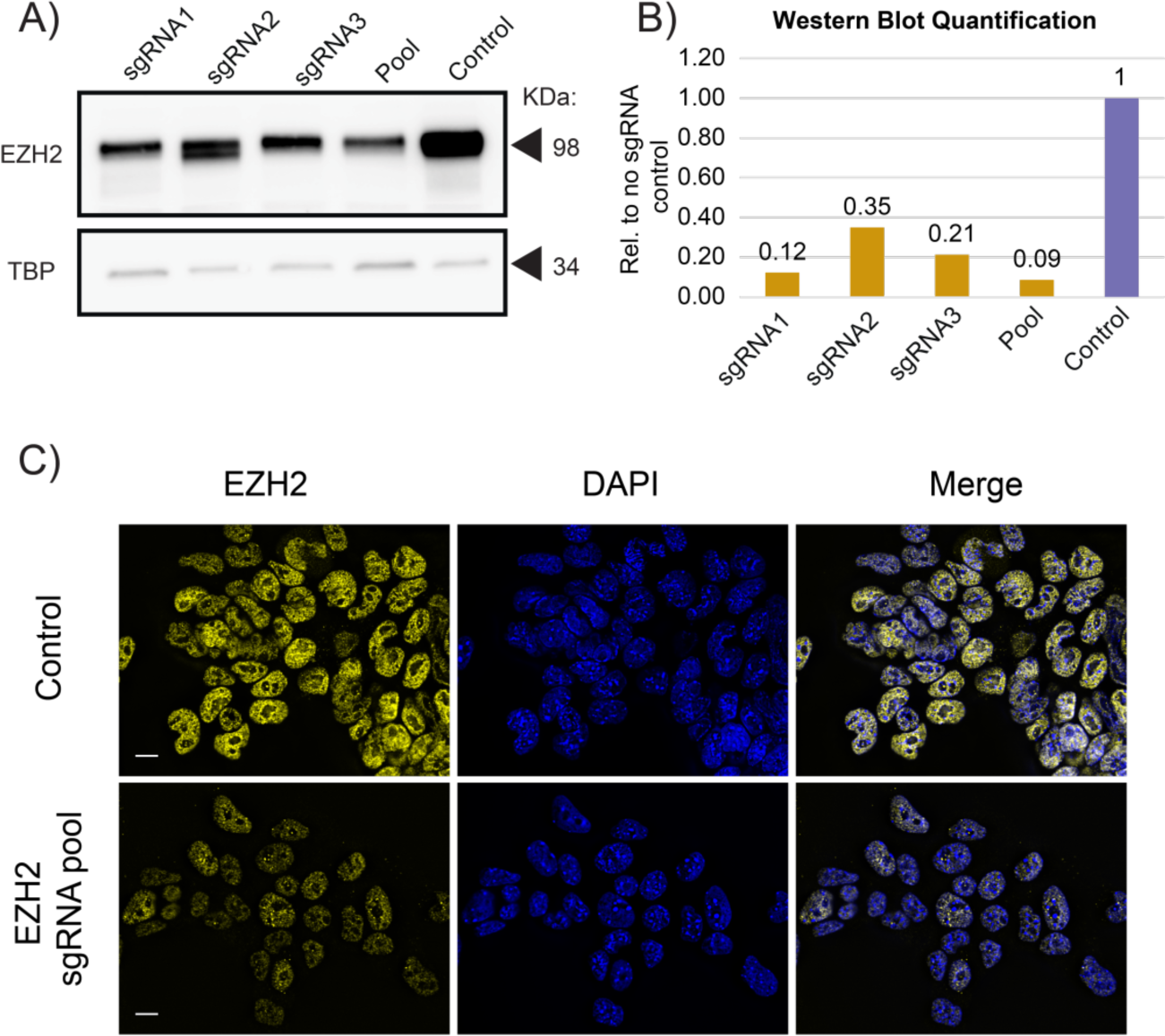
Inducible protein-coding gene knockdown with CRISPR-Bac. **(A)** Representative western blot of CRISPR-Bac assay in which three separate sgRNAs targeting *Ezh2*, a pool of all three sgRNAs, or a non-targeting sgRNA control (“Control”) were co-transfected along with the inducible Cas9-expressing piggyBac cargo at a 2:2:1 rtTA-sgRNA:Cas9:transposase ratio into E14 mouse ESCs and selected with Hygromycin B and G418 for 10 days. Western blots to EZH2 and TBP were performed on protein extracted from stably-selected cells, after four days of Cas9 induction with doxycycline. **(B)** Quantitation of Western blot data from (A). **(C)** Representative immunofluorescence image showing EZH2 knockdown in non-targeting sgRNA control (“control”) or pooled sgRNA cells, each transfected with 8:2:1 rtTA-sgRNA:Cas9:transposase ratio.

### Targeted deletion of genetic elements using CRISPR-Bac

An important use of CRISPR-Cas9 is to create targeted deletions of regulatory elements (Hsu et al., 2014; Wright et al., 2016). To test the utility of CRISPR-Bac in this application, we cloned into CRISPR-Bac pairs of sgRNAs that flank multiple different regulatory elements (RE1, RE2, RE3, RE4), and created ESCs that stably express the different sgRNA pairs along with doxycycline-inducible Cas9. We induced expression of Cas9 for 4 days, collected genomic DNA, and performed quantitative PCR (qPCR) using amplicons within the deleted regions. By comparing qPCR results between the sgRNA-expressing ESCs and non-targeting sgRNA control ESCs, we approximated the extent that each targeted region was deleted in a polyclonal cell population. For 4 of 4 deletions, we observed more than 40% reduction in signal, implying at least one allele was deleted in most cells of the population (Figure 3A).

**Figure 3.**
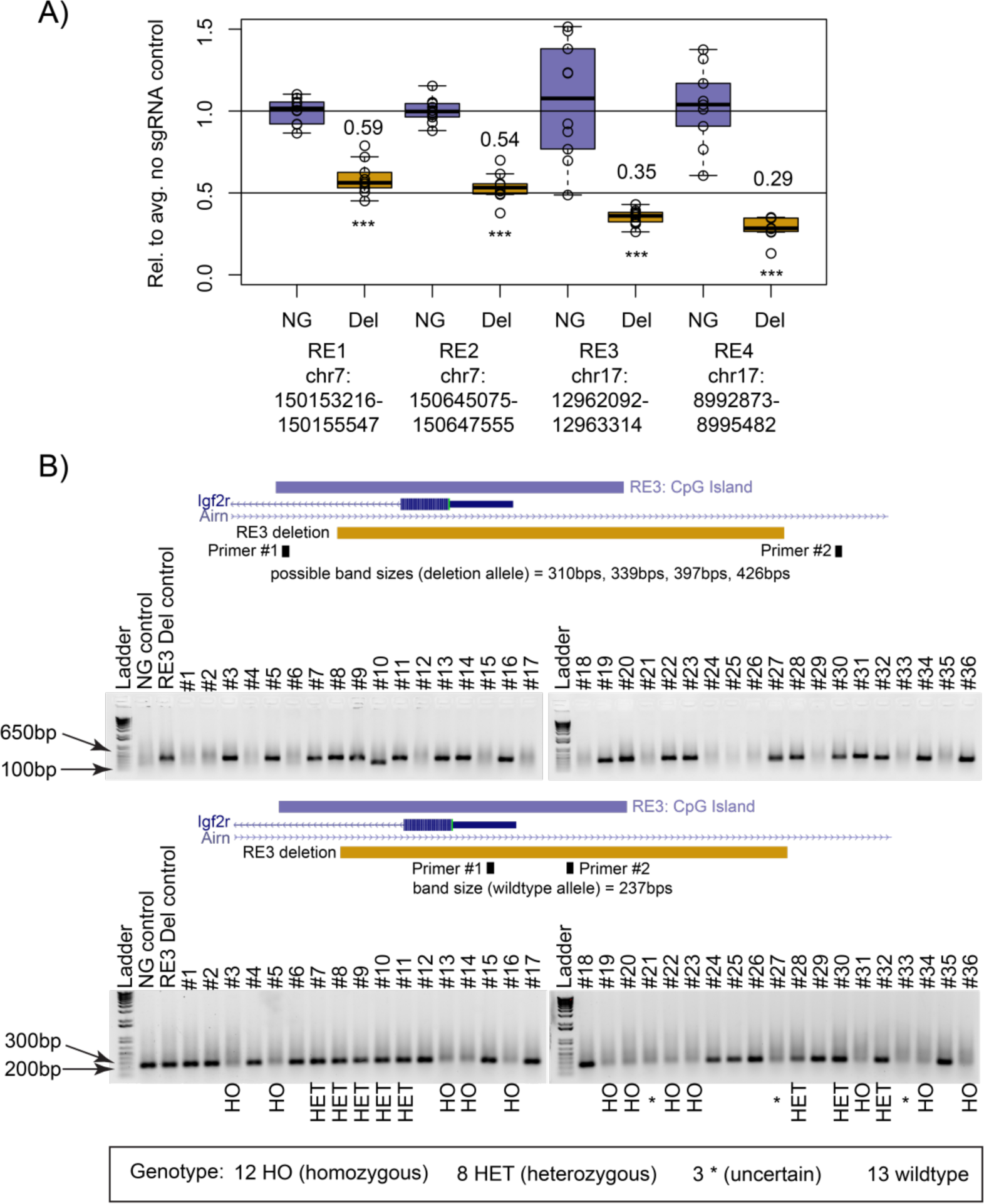
Targeted deletion of DNA regulatory elements using CRISPR-Bac. **(A)** qPCR results from polyclonal populations of ESCs expressing Cas9 and pairs of sgRNAs flanking four separate regulatory elements (RE). Cells were transfected with an 8:2:1 rtTA-sgRNA:Cas9:transposase ratio and Cas9 was induced with doxycycline for 4 days. Primers for qPCR sit entirely inside of the expected deletion. Ten qPCR technical replicates were performed for each assay, and each replicate in the non-targeting sgRNA control (NG) and sgRNA-expressing cells (Del) is shown relative to the average of the signal in NG. ***, p < 0.001 from a two-sided t-test between NG and Del. **(B)** Agarose gel showing genotyping PCR products for the NG control and RE3 Del polyclonal populations from (A) and 36 clones isolated from RE3 deletion ESCs. The UCSC browser tracks above each gel show the location of RE3, the location of the expected deletion, and the location of primers used in the corresponding genotyping PCR. The top gel identifies clones that have a deleted allele (four possible band sizes based on the combination of sgRNAs that cut). The bottom gel identifies clones that have a wildtype allele (primer pairs are the same as used for RE3 in (A)).

As a parallel method to assess the deletion efficiency using CRISPR-Bac, we isolated 36 individual colonies from cells transfected with sgRNAs to the RE3 element, and extracted their genomic DNA. To assess whether a deletion occurred on at least one allele, we performed PCR using primers that flanked the expected deletion. 21 of 36 clones (58%) showed a band within the expected size range (310bp-426bps; Figure 3B, top panel). This signified that these clones were at least heterozygous for the deletion. To distinguish between clones that were heterozygous versus homozygous for the deletion, we used a pair of primers that amplify inside the deletion. 12 out of the 21 clones (33% of the 36 clones) did not show a band, indicating that no wildtype allele was present and the cells were homozygous for the deletion (Figure 3B, bottom panel). These data demonstrate that CRISPR-Bac can be used to generate targeted genomic deletions with high efficiency.

### Activation and repression of protein-coding gene transcription using CRISPR-Bac

In addition to creating targeted genomic deletions, the CRISPR-Cas9 system can be used to up- or down-regulate genes from their endogenous promoters, by targeting dCas9 fused to effector domains that recruit transcriptional co-activators or co-repressors. We cloned one such transcriptional activator fusion, dCas9-VP160 from (Cheng et al., 2013), and one such transcriptional repressor fusion, dCas9-KRAB from (Kearns et al., 2014), into the same piggyBac-based inducible expression vector that we used to express catalytically active Cas9 (Figure 1). We then tested our ability to upregulate *Ascl1*, a silent gene in ESCs, with dCas9-VP160, and we tested our ability to downregulate *Oct4*, an active gene in ESCs, with dCas9-KRAB (Figure 4A). After 2 days of doxycycline treatment, we routinely observed 350-fold upregulation of *Ascl1* relative to non-targeting sgRNA control cells (Figure 4B). Using multiple sets of published and in-house-designed sgRNAs, the maximum level of *Oct4* down regulation we achieved was two- to three-fold (Figure 4C and data not shown). These data show that CRISPR-Bac can be used to up- and down-regulate the transcription of protein-coding genes of interest.

**Figure 4.**
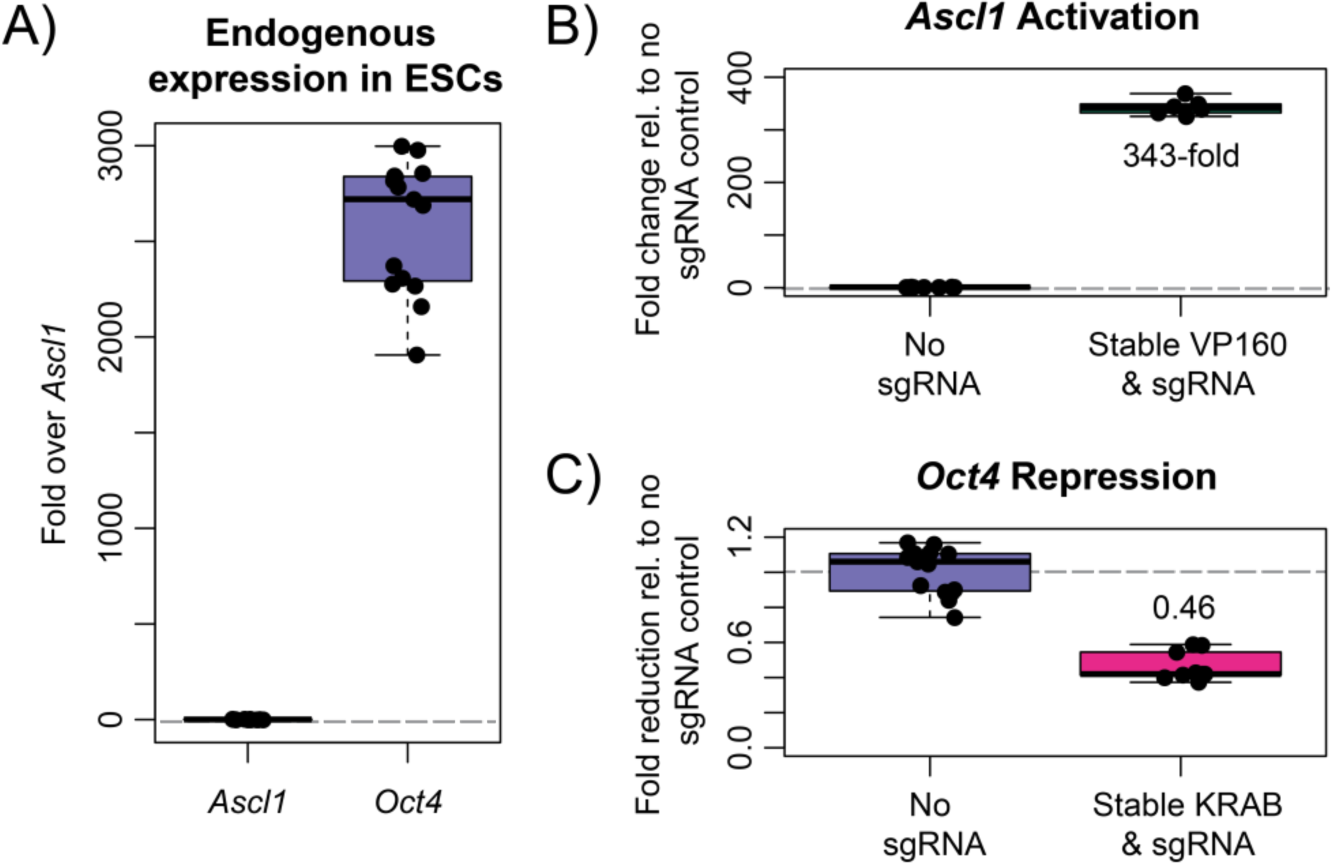
Activation and repression of protein-coding gene transcription using CRISPRBac. All RNA for qPCR analysis was harvested after 2 days of doxycycline-induction of the dCas9 fusion. Each replicate is relative to the average of the signal in the non-targeting sgRNA control. **(A)**qPCR results showing endogenous expression of *Ascl1* and *Oct4* in ESCs relative to the average *Ascl1* signal. **(B)** qPCR showing transcriptional activation of *Ascl1* using dCas9-VP160 and a pool of four *Ascl1*-targeting sgRNAs from (Perez-Pinera et al., 2013) transfected at a 1:1:2 rtTA-sgRNA:dCas9-VP160:transposase ratio. Data points are from 2 biological replicates with 3 qPCR technical replicates each. **(C)** qPCR showing transcriptional repression of *Oct4* using dCas9-KRAB. Data are from 3 independent experiments that used either a 1:1:2, 8:2:1, and 15:5:1 rtTA-sgRNA:dCas9-KRAB:transposase ratio. Two independent sets of sgRNAs were used in separate experiments.

### Activation and repression of lncRNA transcription using CRISPR-Bac

We next examined whether we could use CRISPR-Bac to activate and repress transcription of a lncRNA using dCas9-VP160 and dCas9-KRAB, respectively. We chose to target a lncRNA called *Airn* in two cell types: mouse ESCs, where *Airn* is naturally expressed at low levels, and mouse trophoblast stem cells (TSCs), where *Airn* is expressed and active ((Andergassen et al., 2017; Calabrese et al., 2015; Latos et al., 2009); Figure 5A). In ESCs, we were able to activate *Airn* ~12-fold above its levels in non-targeting sgRNA control cells (Figure 5B), but we were not able to repress *Airn*, likely due to its low endogenous expression (not shown; (Latos et al., 2009)). In TSCs, we were able to repress *Airn* to 10% of its normal expression and activate *Airn* 2.5-fold relative to non-targeting sgRNA control cells (Figure 5C). Therefore, CRISPR-Bac can be used to activate and repress transcription of lncRNAs.

**Figure 5.**
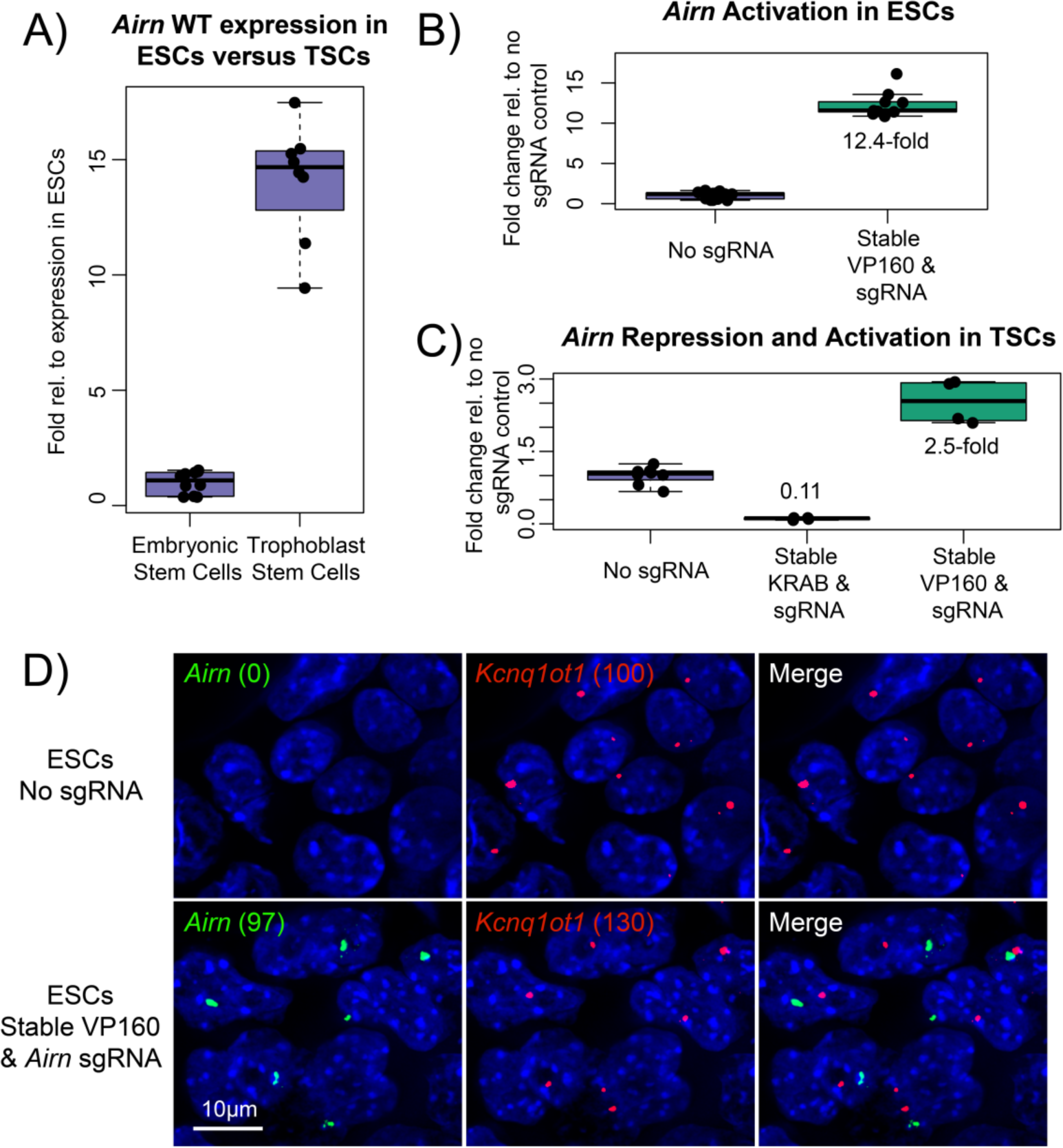
Activation and repression of lncRNA transcription using CRISPR-Bac. **(A)** qPCR showing relative levels of *Airn* expression in wildtype ESCs and TSCs. **(B)** qPCR measuring *Airn* expression in ESCs with stably transfected dCas9-VP160 and *Airn*-targeting sgRNA, two days after dox induction. **(C)** qPCR results for TSCs with stably transfected dCas9-KRAB or dCas9-VP160 and *Airn*-targeting sgRNA, after four days of doxycycline induction. **(D)** Representative RNA FISH image showing *Airn* and *Kcnq1ot1* RNA in ESCs harvested alongside ESCs from (B). Numbers in parenthesis correspond to spots counted by Imaris software for *Airn* and *Kcnq1ot1* in each cell line.

Under normal physiological conditions, the *Airn* lncRNA is monoallelically expressed due to a process called genomic imprinting that leads to methylation of its promoter and gene silencing specifically on the maternally inherited allele (Lee and Bartolomei, 2013; Stoger et al., 1993). To assess whether activation of *Airn* via CRISPR-Bac led to mono- or bi-allelic activation of the lncRNA, we performed RNA Fluorescence In Situ Hybridization (FISH) in ESCs stably expressing dCas9-VP160 and either a non-targeting sgRNA or an *Airn*-targeting sgRNA. We performed a two-color RNA FISH experiment where one probe was complementary to the *Airn* lncRNA, and the other probe was complementary to the *Kcnq1ot1* lncRNA. *Kcnq1ot1*, like *Airn* is also imprinted and monoallelically expressed (Lee and Bartolomei, 2013). Unlike *Airn*, *Kcnq1ot1* it is robustly expressed in ESCs under normal conditions (Umlauf et al., 2004). *Kcnq1ot1* therefore served as a control to gauge the extent of *Airn* monoallelism upon activation by CRISPR-Bac. After taking z-stacks on a widefield microscope and deconvolving the resultant images, we used an automated pipeline to identify puncta whose RNA FISH signal surpassed a specified threshold. In two images taken of cells expressing the non-targeting sgRNA control, we counted zero puncta of *Airn* relative to 100 puncta of *Kcnq1ot1*, confirming prior data that show *Kcnq1ot1* is robustly expressed in ESCs while *Airn* is not (Latos et al., 2009; Umlauf et al., 2004). In contrast, in two images taken of cells expressing the *Airn*-targeting sgRNA, we counted 97 puncta of *Airn* relative to 130 puncta of *Kcnq1ot1* (Figure 5D). These data support the notion that CRISPR-Bac activates expression of *Airn* on the unmethylated paternal allele, and that the methylated maternal allele of *Airn* remains resistant to activation (Stoger et al., 1993).

### sgRNA titration to achieve variable levels of lncRNA induction

The number of piggyBac cargos inserted into the genome can be controlled by altering the ratio of cargo vector to transposase plasmid (Cadinanos and Bradley, 2007; Wang et al., 2008; Wilson et al., 2007). The CRISPR-Bac platform relies on simultaneous delivery of two cargo vectors: one vector expressing the sgRNA and rtTA/G418 resistance genes, and the other vector expressing the Cas9/dCas9 variant and hygromycin resistance genes (Figure 1). We sought to determine whether the extent of activation of a target gene of interest could be altered by altering the ratios of sgRNA, Cas9, and piggyBac transposase vectors in transfections. We tested a range of sgRNA-to-dCas9VP160-to-transposase ratios, using the *Airn* lncRNA as our target gene for activation (Table S2). We found modest but significant differences in the level of *Airn* activation when we transfected higher amounts of sgRNA and dCas9-VP160 plasmids relative to the piggyBac transposase plasmid (Figure 6A; see table of adjusted p-values from Tukey’s HSD test), and these differences were accompanied by increased numbers of sgRNA and dCas9-VP160 cargo insertions per cell (Figure 6B). Thus, the extent of target gene activation using CRISPR-Bac can be partly controlled by changing the ratios of sgRNA/rtTA, Cas9, and transposase plasmids in transfections.

**Figure 6.**
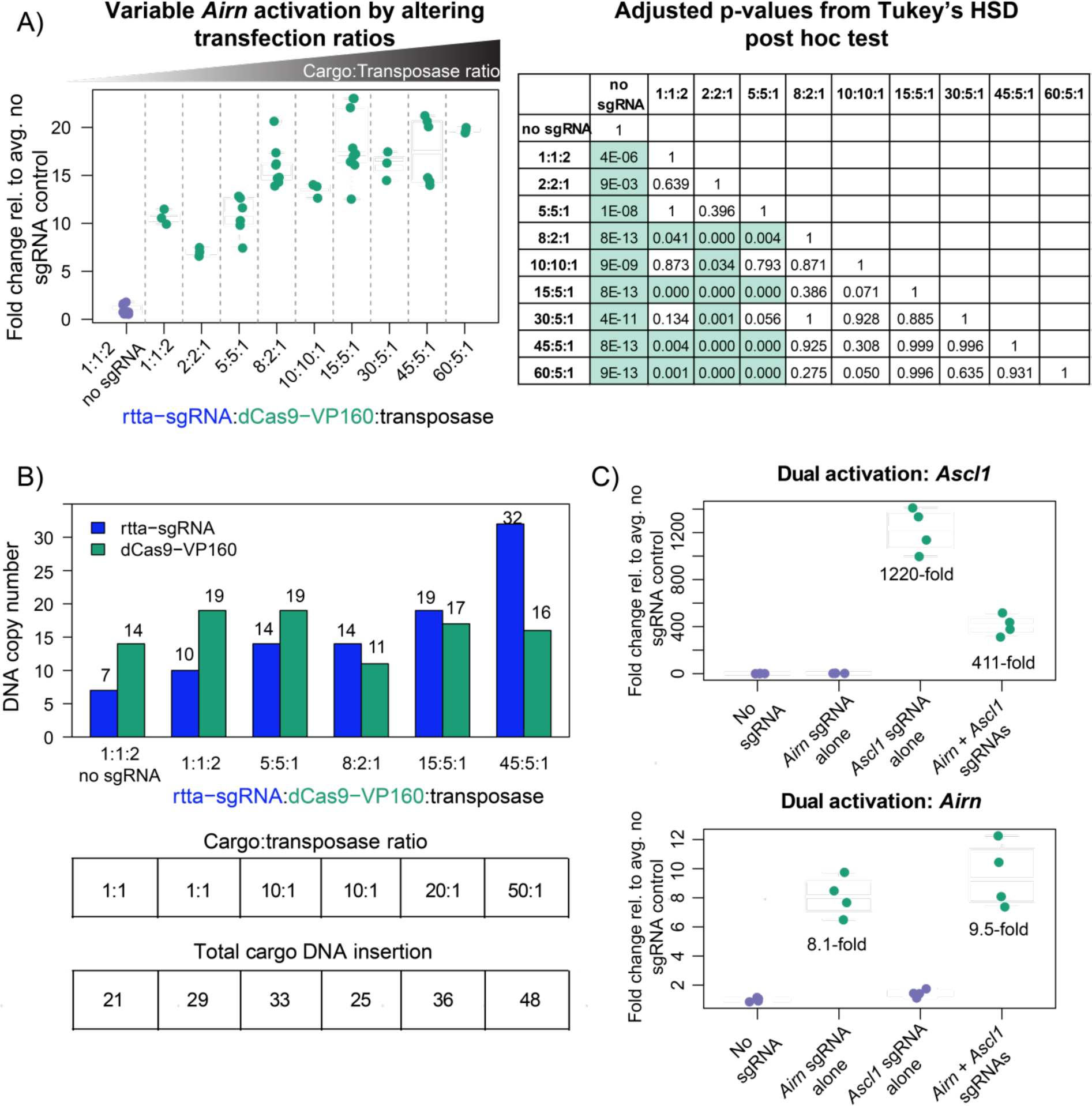
Cargo to transposase ratio controls the extent of activation and multiplex gene activation by CRISPR-Bac. RNA for all experiments was harvested two days after dox induction. Each qPCR replicate is shown relative to the average of the signal in cells transduced with our non-targeting sgRNA control (“no sgRNA”). **(A)** qPCR measuring *Airn* activation. X-axis gives the transfection ratio of rtTA-sgRNA to dCas9-VP160 to transposase for each experiment. Ratios are plotted in ascending order based on the summed cargo (rtTA-sgRNA plus dCas9-VP160) to transposase ratio. “1:1:2 no sgRNA” and “8:2:1” data are the same as shown in Figure 5B. For each experiment, data points are from at least one biological replicate with three qPCR technical replicates. Corresponding table gives adjusted p-values from Tukey’s HSD post hoc test for all comparisons, where p <= 0.05 are highlighted in green. **(B)** Bar plot showing DNA copy number per cell for the rtTA-sgRNA and dCas9-VP160 cargos under each transfection condition. Numbers over each bar give the average copy number calculated from 3 technical qPCR replicates. The tables below correspond to the bar plot showing the transfection ratio and total number of DNA cargos inserted (copy number of rtTA-sgRNA plus dCas9-VP160). **(C)** Simultaneous activation of *Ascl1* and *Airn* transcription upon co-transfection of a pool of four *Ascl1* sgRNAs from (Perez-Pinera et al., 2013) and one *Airn* sgRNA at an 8:2:1 (rtTA-sgRNA:dCas9-VP160:transposase) ratio. Data points are from two biological replicates with two qPCR technical replicates each.

### Simultaneous upregulation of two genes via CRISPR-Bac

By co-expression of multiple sgRNAs, CRISPR can be used to activate or repress multiple genes simultaneously (Cheng et al., 2013). To test if CRISPR-Bac is capable of multiplexed gene activation, we created ESCs expressing dCas9-VP160 and sgRNAs targeting the *Ascl1* and *Airn* promoters (same sgRNAs as in Figures 4B, 5B-C, and 6A). Relative to non-targeting sgRNA controls, qPCR demonstrated simultaneous 411-fold activation of *Ascl1* and 9.5-fold activation of *Airn* when sgRNAs for both targets were co-transfected (Figure 6C). This confirms that CRISPR-Bac can be used to target multiple genes in a single experiment.

## Conclusions

In *Mus musculus*-derived embryonic stem cells and trophoblast stem cells, we have shown that CRISPR-Bac can be used to knockdown proteins through frameshift/deletion, to delete kilobase-sized regulatory elements with high efficiency, and to up- and down-regulate the transcription of protein-coding genes and an imprinted lncRNA. Levels of CRISPR-induced activation could partly be controlled through delivery of different ratios of CRISPR-Bac vectors. It seems likely that the use of different promoter elements within CRISPR-Bac (for example, a constitutive CMV promoter driving dCas9-VP160 instead of a TRE) might afford additional levels of control. It may also be possible to engineer CRISPR-Bac vectors that express multiple sgRNAs, as has been done elsewhere (Albers et al., 2015; Kabadi et al., 2014; Sakuma et al., 2014). Although in this work we only tested CRISPR-Bac in two cell types derived from *Mus musculus*, it seems reasonable to presume that the CRISPR-Bac vectors we described or their modified derivatives would be functional in other mammalian cell types, given the broad activity of the piggyBac transposase (Cadinanos and Bradley, 2007; Ding et al., 2005; Kahlig et al., 2010; Kaji et al., 2009; Li et al., 2011; Li et al., 2013; Wang et al., 2008; Wilson et al., 2007; Yusa et al., 2009). By circumventing the need to package CRISPR-Cas9 components into lentiviral delivery systems, CRISPR-Bac provides a convenient and versatile platform for inducible genome editing.

## Materials and Methods

### Construction of CRISPR-Bac vectors

To create the doxycycline-inducible Cas9, dCas9-VP160, and dCas9-KRAB piggyBac vectors, a parent piggyBac vector was created in which a bGH-polyA signal and an EF1a promoter driving expression of a hygromycin resistance gene was ligated into the cumate-inducible piggyBac transposon vector from System Biosciences after its digestion with HpaI and SpeI, which cut just downstream of each chicken b-globin insulator sequence and removed all other internal components of the original vector. The TRE from pTRE-Tight (Clontech) was cloned upstream of the bGH-polyA site, and Cas9 from pX330 (Addgene plasmid # 42230; (Cong et al., 2013); gift from Feng Zhang), dCas9-VP160 from (Addgene plasmid # 48225; (Cheng et al., 2013); gift from Rudolf Jaenisch) and dCas9-KRAB from (Addgene plasmid # 50917;(Kearns et al., 2014); gift from Rene Maehr & Scot Wolfe) were each cloned behind the TRE by digestion with AgeI and SalI (NEB) followed by Gibson Assembly (NEB), to generate piggyBac cargo vectors capable of inducibly expressing Cas9, dCas9-VP160, and dCas9-KRAB, respectively, upon addition of doxycycline.

To create the rtTA-sgRNA expressing piggyBac vector, the dual BbsI sites in pX330 were converted to BsmbI sites using oligonucleotides, and the entire U6 expression cassette was cloned via Gibson assembly into the PacI site upstream of the rtTA3-IRES-Neo cassette in the rtTA-piggyBac-Cargo vector described in (Kirk et al., 2018). The rtTA3-IRES-Neo cassette was originally cloned from pSLIK-Neo and was a gift from Iain Fraser (Addgene plasmid # 25735). Oligonucleotides used for cloning are in Table S1.

### sgRNA Design

Oligonucleotides used for sgRNA cloning are listed in Table S1. Guide RNAs were designed using the CRISPOR program or taken from published sources ((Haeussler et al., 2016); Table S1).

### Embryonic stem cell (ESC) culture

ESCs were grown on gelatin coated plates at 37°C in a humidified incubator at 5% CO2. Media was changed daily and consisted of DMEM high glucose plus sodium pyruvate, 0.1 mM non-essential AA, 100 u/mL penicillin-streptomycin, 2 mM L-glutamine, 0.1mM 2-mercaptoethanol, 15% ES-qualified FBS, and 1:500 LIF conditioned media produced from Lif—1Cα (COS) cells. ESCs were split at an approximate ratio of 1:6 every 48hr.

### Trophoblast stem cell (TSC) culture

TSCs were cultured as in (Quinn et al., 2006). Briefly, TSCs were cultured at 37°C on pre-plated irradiated MEF feeder cells in TSC media [RPMI (Invitrogen), 20% Qualified FBS (Invitrogen), 100 u/mL penicillin-streptomycin, 1mM sodium pyruvate (Invitrogen), 100µM β-mercaptoethanol (Sigma), and 2mM L-glutamine] supplemented with Fgf4 (25ng/ml; Invitrogen) and Heparin (1µg/ml; Sigma) just before use. At passage, TSCs were trypsinized with 0.125% Trypsin (Invitrogen) for 3 minutes at room temperature and gently dislodged from their plate with a sterile, cotton-plugged Pasteur pipette (Thermofisher). To deplete MEF feeder cells from TSCs prior to RNA isolation, TSCs were pre-plated for 40 minutes and cultured for three days in 70% MEF-conditioned TSC media supplemented with Fgf4 (25ng/ml; Invitrogen) and Heparin (1µg/ml; Sigma).

### Transfections

To generate stable CRISPR-Bac cell lines, 5×10^5 E14 embryonic stem cells were seeded in a single well of a 6-well plate, and the next day transfected with piggyBac cargo vectors and pUC19-piggyBac transposase from (Kirk et al., 2018), totaling 2.5 µg of plasmid DNA (see exact amounts in Table S2), using Lipofectamine 3000 (Invitrogen) according to manufacturer instructions. Cells were subsequently selected on Hygromycin [150µg/ml; Gibco] and G418 [200µg/ml; Gibco] for 7 to 12 days. Due to the efficiency of piggyBac cargo integration and the rapidity of Hygromycin selection, most observable death from drug selection occurred within ~3 days after addition of Hygromycin and G418 (i.e. cells with Hygromycin resistance were invariably resistant to G418).

### Protein isolation and western blotting

To isolate protein for western blotting, ESCs were washed with PBS, and then lysed with RIPA buffer (10 mM Tris-Cl (pH 7.5), 1 mM EDTA, 0.5 mM EGTA, 1% NP40, 0.1% sodium deoxycholate, 0.1% SDS, 140 mM NaCl) supplemented with 1 mM PMSF (Fisherscientific) and 1x protease inhibitor cocktail (Sigma) for 15 minutes at 4C, four days after induction with 1µg/ml doxycycline. Prior to western blotting, protein levels were quantified using the DC assay from Biorad. For western blotting, primary and secondary antibody incubations were done for 1hr at room temperature. Antibodies used were EZH2 (Cell Signaling #5246, 1:1000 dilution), TBP (Abcam ab818, 1:2000 dilution), donkey anti-mouse IgG-HRP secondary (Santa Cruz; sc-2314; 1:2500), and donkey anti-rabbit IgG-HRP secondary (Santa Cruz; sc-2313; 1:2500).

### Genomic DNA isolation and qPCR

To isolate genomic DNA, 400ul of ESC lysis buffer (100 mM Tris-HCl, pH 8.1, 5mM EDTA, pH 8.0, 200mM NaCl, 0.2% SDS) supplemented with 80ul proteinase K (Denville) was used per 24-well well of ESCs, four days after induction with 1µg/ml doxycycline. Lysed ESCs were incubated at 55°C overnight, cells were boiled at 100°C for 1 hr to degrade RNA, and DNA was precipitated by addition of 2 volumes of 100% ethanol. DNA was pelleted and resuspended in 1x TE (10mM Tris-HCl 1mM EDTA pH 8.0) overnight at room temperature prior to qPCR. qPCR was performed using 100 ng of DNA per reaction and iTaq Universal SYBR Green Supermix (Biorad), with primers specified in Table S1. All related plots were generated using R version 3.4.1 (Team, 2017).

### qPCR for DNA copy number analysis

Genomic DNA was prepared as in Genomic DNA isolation and qPCR section above. qPCR signal (SsoFast, Biorad) from the genomic DNA was compared to signal from a molar standard amplified from increasing amounts of the corresponding dCas9-VP160 and rtTA plasmids. Primers used are listed in Table S1. All related plots were generated using R version 3.4.1 (Team, 2017).

### Generation of clonal ESCs with targeted genomic deletions and genotyping

After 4 days of dox induction, RE3 deletion E14 cells were cultured two days in the absence of dox to ensure that Cas9 was fully depleted. Then, 2,000 cells were plated on a 10cm plate with pre-plated irradiated MEF feeder cells. After 4 days, individual colonies were picked and plated on irMEFs. Clonal lines were passaged twice off of MEFs before genomic DNA was prepared as in Genomic DNA isolation and qPCR section above.

Genotyping PCR reactions were performed with gDNA using Apex Taq DNA Polymerase (Genesee Scientific). The first set of primers flanked the deletion and identified clonal lines with at least one allele deleted. The second set only amplified a wildtype allele, with both primers sitting inside the deletion. Many clones showed weak wildtype bands, likely due to MEF gDNA and not due to the presence of a wildtype allele in the ESC clone. Primers used are listed in Table S1.

### RNA Isolation and qPCR

RNA was isolated using Trizol (Invitrogen). For RT-qPCR assays, 2µg of RNA was reverse transcribed using MultiScribe RT (Applied Biosystems), and qPCR was performed using iTaq Universal SYBR Green (Biorad) and primers specified in Table S1. All related plots were generated using R version 3.4.1 (Team, 2017).

### RNA FISH

Fosmid Wl1-2156F18 (*Airn*) and BAC RP23-101N20 (*Kcnq1ot1*) were ordered from the BACPAC resource center and fingerprinted with restriction digestion prior to use to verify inserted DNA. Fluorescent labeling was performed using BioPrime (Invitrogen). ESCs were fixed on coverslips for 10 minutes in 4% paraformaldehyde/PBS, followed by a 10-minute permeabilization on ice in 0.5% TritonX-100 in PBS and 1:200 Ribonucleoside Vanadyl Complex (NEB). Coverslips were stored at -20C in 70% ethanol until use.

To initiate the RNA FISH protocol, coverslips were dehydrated by serial 3-minute incubations with 75%, 85%, 95%, and 100% ethanol, and air-dried for 5 minutes. RNA FISH probes were added and coverslips were placed cell-side down in a chamber humidified with 50% formamide/2xSSC overnight at 37°C. After overnight incubation, coverslips were washed 3x with 50% formamide/2xSSC at 42C and 3x with 1xSSC at 50C. Each wash was 5 minutes long. Coverslips were then rinsed 1x with PBS before a 2 minute incubation in DAPI stock diluted 1:1000 in water. Coverslips were rinsed twice more and affixed to glass slides using Vectashield (VectorLabs), then sealed with nail polish.

Four dimensional datasets were acquired by taking multi-channel Z-stacks on an Olympus BX61 widefield fluorescence microscope using a Plan-Aprochromat 63X/1.4 oil objective and a Hamamatsu ORCA R2 camera, controlled by Volocity 6.3 software. Excitation was provided by a mercury lamp and the following filters were used for the three fluorescent channels that were imaged: 377/25 ex, 447/30 em for DAPI (DAPI-5060B Semrock filter); 482/17 ex, 536/20 em for AlexaFluor488 (Semrock FITC-3540B filter); 562/20 ex, 642/20 em for Cy3 (Semrock TXRED-4040B filter). Pixel size was 0.108 µm, Z spacing was 0.2 µm, and images had 1344×1024 pixels. Between 46-49 Z-stacks were acquired for each image. Z-stacks were deconvolved using the iterative-constrained algorithm (Mediacy AutoQuantX3) with default algorithm settings. Sample settings for the deconvolution were: peak emissions for dyes (570 nm, 519 nm, 461 nm for Cy3, AlexaFluor 488 and DAPI respectively), widefield microscopy mode, NA = 1.4, RI of oil = 1.518, and RI of sample = 1.45. After deconvolution, RNA FISH signals were located using the “Spots” function in Imaris software (version 8.3.1) and marked with equal sized spheres. To initially call spots on all images, spot detection values were set at 0.5µm for xy and 1.5µm for z, and background subtraction and auto quality settings were used. We manually optimized the quality/sensitivity setting to call *Kcnq1ot1* spots, and then used the same quality threshold to call *Airn* spots for the same image. Images are shown as maximum intensity projections made using ImageJ (Schindelin et al., 2012).

### Immunofluorescence (IF)

ESCs were fixed on coverslips the same as for RNA FISH (see above). To initiate the IF protocol, coverslips were washed twice in PBS and blocked for 30 minutes at room temperature in blocking solution (1x PBS with 0.2% Triton X-100, 1% goat serum, and 6 mg/mL IgG-free BSA). Then, coverslips were washed in 0.2% triton/1x PBS and incubated with EZH2 antibody (Cell Signaling #5246; 1:200 in block solution) for 1 hour at RT. Coverslips were washed 3x in 0.2% triton/1x PBS for 4 minutes each and incubated with secondary antibody (AlexaFluor 647 goat anti-rabbit, A-21245, 1:1000 in block solution) for 30 minutes at RT. After incubation, coverslips were washed 3x in 0.2% triton/1x PBS for 4 minutes each and rinsed 1x with PBS before a 2 minute incubation in DAPI stock diluted to 5ng/ml in water. Coverslips were rinsed twice more and mounted to glass slides using Prolong Gold (Thermo Fisher Scientific P10144). Imaging and deconvolution was performed the same as described in the RNA FISH section with the below exceptions. The filters used for the two fluorescent channels that were imaged are 377/25 ex, 447/30 em for DAPI (DAPI-5060B Semrock filter) and 628/20 ex, 692/20 em for AlexaFluor 647 (Semrock Cy5 4040A filter). Approximately 40 Z-stacks were acquired for each image. Sample settings for the deconvolution included the following peak emissions for dyes: 670 nm and 461 nm for AlexaFluor 647 and DAPI, respectively. Images are shown as maximum intensity projections made using ImageJ (Schindelin et al., 2012).

## Acknowledgments

We thank UNC colleagues for discussions. This work was supported by National Institutes of Health (NIH) Grant GM121806, Basil O’Connor Award #5100683 from the March of Dimes Foundation, and funds from the Lineberger Comprehensive Cancer Center and UNC Department of Pharmacology (J.M.C.). E.T., D.M.L., E.R.H. were supported in part by the National Institute of General Medical Sciences under award T32 GM007092. K.C.A.B. and R.M.M. were supported in part by the NIGMS training award T32 GM119999. E.C.M.V. was supported in part by the NIEHS Toxicology Training Grant T32 ES007126. The Microscopy Services Laboratory was supported in part by NCI grant P30 CA016086.

## Author Contributions

M.D.S. and J.M.C. conceived the study, M.D.S., E.T., K.C.A.B., D.M.L., E.R.H., R.M.M., S.O.K., and E.C.M.V. performed the experiments, M.D.S. and J.M.C. performed the analysis, and M.D.S. and J.M.C. wrote the paper.

## Declaration of interests

The authors declare no competing interests.

**Table S1. Oligonucleotides used.**Related to all figures. Table has 3 columns, including the name, sequence and usage of all oligonucleotides in the paper. ** in front of the name indicates sgRNAs from published papers [(Kearns et al., 2015) for Oct4 and (Perez-Pinera et al., 2013) for Ascl1].

**Table S2. DNA transfection ratios.**Related to Figure 6. For each ratio (rtTA-sgRNA:dCas9-VP160:transposase) in Figure 6, this table gives the amount of transfected DNA (in nanograms) for each individual plasmid.

## References

Albers, J., Danzer, C., Rechsteiner, M., Lehmann, H., Brandt, L.P., Hejhal, T., Catalano, A., Busenhart, P., Goncalves, A.F., Brandt, S., et al. (2015). A versatile modular vector system for rapid combinatorial mammalian genetics. J Clin Invest 125, 1603–1619.

Andergassen, D., Dotter, C.P., Wenzel, D., Sigl, V., Bammer, P.C., Muckenhuber, M., Mayer, D., Kulinski, T.M., Theussl, H.C., Penninger, J.M., et al. (2017). Mapping the mouse Allelome reveals tissue-specific regulation of allelic expression. eLife 6.

Cadinanos, J., and Bradley, A. (2007). Generation of an inducible and optimized piggyBac transposon system. Nucleic Acids Res 35, e87.

Calabrese, J.M., Starmer, J., Schertzer, M.D., Yee, D., and Magnuson, T. (2015). A survey of imprinted gene expression in mouse trophoblast stem cells. G3 (Bethesda) 5, 751–759.

Cheng, A.W., Wang, H., Yang, H., Shi, L., Katz, Y., Theunissen, T.W., Rangarajan, S., Shivalila, C.S., Dadon, D.B., and Jaenisch, R. (2013). Multiplexed activation of endogenous genes by CRISPR-on, an RNA-guided transcriptional activator system. Cell Res 23, 1163–1171.

Cong, L., Ran, F.A., Cox, D., Lin, S., Barretto, R., Habib, N., Hsu, P.D., Wu, X., Jiang, W., Marraffini, L.A., et al. (2013). Multiplex genome engineering using CRISPR/Cas systems. Science 339, 819–823.

Ding, S., Wu, X., Li, G., Han, M., Zhuang, Y., and Xu, T. (2005). Efficient transposition of the piggyBac (PB) transposon in mammalian cells and mice. Cell 122, 473–483.

Gossen, M., Freundlieb, S., Bender, G., Muller, G., Hillen, W., and Bujard, H. (1995). Transcriptional Activation by Tetracyclines in Mammalian-Cells. Science 268, 1766–1769.

Haeussler, M., Schonig, K., Eckert, H., Eschstruth, A., Mianne, J., Renaud, J.B., Schneider-Maunoury, S., Shkumatava, A., Teboul, L., Kent, J., et al. (2016). Evaluation of off-target and on-target scoring algorithms and integration into the guide RNA selection tool CRISPOR. Genome Biol 17, 148.

Hartenian, E., and Doench, J.G. (2015). Genetic screens and functional genomics using CRISPR/Cas9 technology. FEBS J 282, 1383–1393.

Hsu, P.D., Lander, E.S., and Zhang, F. (2014). Development and Applications of CRISPR-Cas9 for Genome Engineering. Cell 157, 1262–1278.

Joung, J., Konermann, S., Gootenberg, J.S., Abudayyeh, O.O., Platt, R.J., Brigham, M.D., Sanjana, N.E., and Zhang, F. (2017). Genome-scale CRISPR-Cas9 knockout and transcriptional activation screening. Nat Protoc 12, 828–863.

Kabadi, A.M., Ousterout, D.G., Hilton, I.B., and Gersbach, C.A. (2014). Multiplex CRISPR/Cas9-based genome engineering from a single lentiviral vector. Nucleic Acids Res 42, e147.

Kahlig, K.M., Saridey, S.K., Kaja, A., Daniels, M.A., George, A.L., Jr., and Wilson, M.H. (2010). Multiplexed transposon-mediated stable gene transfer in human cells. Proc Natl Acad Sci U S A 107, 1343–1348.

Kaji, K., Norrby, K., Paca, A., Mileikovsky, M., Mohseni, P., and Woltjen, K. (2009). Virus-free induction of pluripotency and subsequent excision of reprogramming factors. Nature 458, 771–775.

Kearns, N.A., Genga, R.M., Enuameh, M.S., Garber, M., Wolfe, S.A., and Maehr, R. (2014). Cas9 effector-mediated regulation of transcription and differentiation in human pluripotent stem cells. Development 141, 219–223.

Kearns, N.A., Pham, H., Tabak, B., Genga, R.M., Silverstein, N.J., Garber, M., and Maehr, R. (2015). Functional annotation of native enhancers with a Cas9-histone demethylase fusion. Nat Methods 12, 401–403.

Kirk, J.M., Kim, S.O., Inoue, K., Smola, M.J., Lee, D.M., Schertzer, M.D., Wooten, J.S., Baker, A.R., Sprague, D., Collins, D.W., et al. (2018). Functional classification of long non-coding RNAs by k-mer content. Nat Genet.

Latos, P.A., Stricker, S.H., Steenpass, L., Pauler, F.M., Huang, R., Senergin, B.H., Regha, K., Koerner, M.V., Warczok, K.E., Unger, C., et al. (2009). An in vitro ES cell imprinting model shows that imprinted expression of the Igf2r gene arises from an allele-specific expression bias. Development 136, 437–448.

Lee, J.T., and Bartolomei, M.S. (2013). X-inactivation, imprinting, and long noncoding RNAs in health and disease. Cell 152, 1308–1323.

Li, M.A., Turner, D.J., Ning, Z., Yusa, K., Liang, Q., Eckert, S., Rad, L., Fitzgerald, T.W., Craig, N.L., and Bradley, A. (2011). Mobilization of giant piggyBac transposons in the mouse genome. Nucleic Acids Res 39, e148.

Li, Z., Michael, I.P., Zhou, D., Nagy, A., and Rini, J.M. (2013). Simple piggyBac transposon-based mammalian cell expression system for inducible protein production. Proc Natl Acad Sci U S A 110, 5004–5009.

Perez-Pinera, P., Kocak, D.D., Vockley, C.M., Adler, A.F., Kabadi, A.M., Polstein, L.R., Thakore, P.I., Glass, K.A., Ousterout, D.G., Leong, K.W., et al. (2013). RNA-guided gene activation by CRISPR-Cas9-based transcription factors. Nat Methods 10, 973–976.

Quinn, J., Kunath, T., and Rossant, J. (2006). Mouse trophoblast stem cells. Methods Mol Med 121, 125–148.

Sakuma, T., Nishikawa, A., Kume, S., Chayama, K., and Yamamoto, T. (2014). Multiplex genome engineering in human cells using all-in-one CRISPR/Cas9 vector system. Sci Rep 4, 5400.

Schindelin, J., Arganda-Carreras, I., Frise, E., Kaynig, V., Longair, M., Pietzsch, T., Preibisch, S., Rueden, C., Saalfeld, S., Schmid, B., et al. (2012). Fiji: an open-source platform for biological-image analysis. Nat Methods 9, 676–682.

Shin, K.J., Wall, E.A., Zavzavadjian, J.R., Santat, L.A., Liu, J., Hwang, J.I., Rebres, R., Roach, T., Seaman, W., Simon, M.I., et al. (2006). A single lentiviral vector platform for microRNA-based conditional RNA interference and coordinated transgene expression. Proc Natl Acad Sci U S A 103, 13759–13764.

Stoger, R., Kubicka, P., Liu, C.G., Kafri, T., Razin, A., Cedar, H., and Barlow, D.P. (1993). Maternal-specific methylation of the imprinted mouse Igf2r locus identifies the expressed locus as carrying the imprinting signal. Cell 73, 61–71.

Team, R.C. (2017). R: A language and environment for statistical computing. (Vienna, Austria: R Foundation for Statistical Computing).

Umlauf, D., Goto, Y., Cao, R., Cerqueira, F., Wagschal, A., Zhang, Y., and Feil, R. (2004). Imprinting along the Kcnq1 domain on mouse chromosome 7 involves repressive histone methylation and recruitment of Polycomb group complexes. Nat Genet 36, 1296–1300.

Wang, W., Lin, C., Lu, D., Ning, Z., Cox, T., Melvin, D., Wang, X., Bradley, A., and Liu, P. (2008). Chromosomal transposition of PiggyBac in mouse embryonic stem cells. Proc Natl Acad Sci U S A 105, 9290–9295.

Wilson, M.H., Coates, C.J., and George, A.L., Jr. (2007). PiggyBac transposon-mediated gene transfer in human cells. Mol Ther 15, 139–145.

Wright, A.V., Nunez, J.K., and Doudna, J.A. (2016). Biology and Applications of CRISPR Systems: Harnessing Nature’s Toolbox for Genome Engineering. Cell 164, 29–44.

Yusa, K., Rad, R., Takeda, J., and Bradley, A. (2009). Generation of transgene-free induced pluripotent mouse stem cells by the piggyBac transposon. Nat Methods 6, 363–369.

